# Proteome-Wide Compositional Signatures of Extremophilic Adaptation

**DOI:** 10.64898/2026.11.23.768582

**Authors:** Kayra Güvenç Duran, Mahamed Salah

## Abstract

Life survives in extreme environments: hydrothermal vents, hypersaline lakes, and polar ice. The proteins of organisms inhabiting these environments remain stable and functional under stress. Such evolutionary adaptations may leave signatures detectable from amino acid composition, without the need for structural analysis. This study compares proteome-wide compositional metrics across 40 prokaryotic organisms, subdivided equally into 4 groups: thermophiles, halophiles, psychrophiles, and mesophiles for control. For each organism, a reference proteome was retrieved from UniProt, and six metrics were computed for every protein and averaged per organism. These metrics included mean protein length, percentage of acidic residues, percentage of basic residues, acidic-to-basic ratio, mean hydrophobicity, and predicted isoelectric point. Groups were compared statistically by either a one-way ANOVA or Kruskal-Wallis test. Followed by Tukey’s or Dunn’s post-hoc tests with a Bonferroni-corrected P threshold. Four metrics, that all measured protein charge, differed significantly between groups. We have found thermophiles were enriched in basic residues relative to every other group. Conversely, halophiles had the most acidic proteomes, differing from all other groups in predicted pI. Neither protein length nor hydrophobicity differed significantly between groups. Psychrophiles were statistically indistinguishable from mesophiles on all six metrics. Furthermore, subdividing halophiles by salt strategy revealed that the acidic signal was confined to salt-in organisms having more than double the variance explained for the acidic-to-basic ratio when compared to salt-out organisms. These findings indicate that environmental adaptation is detectable proteome-wide mainly through charge, not hydrophobicity. Additionally, point to survival strategy predicting composition more reliably than the environment for halophiles.

## INTRODUCTION

High salinity and extreme temperatures can compromise protein dynamics and catalytic activity. Extremophiles are organisms that thrive under these physicochemical conditions. Their proteins are adapted to function optimally in such environments, and they are found across all three domains of life.^1,2^

A proteome is the complete set of proteins encoded by an organism.^3^ Adaptations affecting many proteins can change the amino acid composition and physicochemical properties across a proteome. These adaptations arise in response to evolutionary pressures like the need for proper protein folding across different environments.^4^ Therefore, extremophiles are often classified according to the environmental conditions they thrive in. These classes are not mutually exclusive since organisms can tolerate more than one extreme.

Such classes include halophiles, thermophiles, and psychrophiles. They are often defined to grow optimally in salt solutions, in elevated temperatures, and in cold temperatures respectively. Other classes also exist, such as hyperthermophiles that live in > 80°C, alkaliphiles with a growth optimum at pH > 8, and more. In contrast, mesophiles are often classified as organisms inhabiting moderate environments.^5,6,7^ Mesophiles serve as the comparison group for this study. Comparing the extremophilic groups against mesophiles allows the assessment of compositional signatures of adaptation across environmental conditions.

Prior studies have reported that thermal adaptation contains a shift towards strong hydrophobic amino acids (IPV) and charged residues of both signs.^8^ In contrast, halophilic adaptation has been reported to contain an overrepresentation of acidic residues and a low hydrophobicity.^9^ Cold adaptation also shows a distinct signature with an increase in the frequency of individual amino acids like A, D, S, and T, and a decrease in basic as well as hydrophilic residues.^10^ It is important to note that these are general trends and they are not necessarily true for every organism in the group.

Interestingly, halophiles use two osmotic strategies to thrive in salt solutions, salt-in and salt-out. The salt-in strategy is based on the intracellular storage of salts including KCl over 37% (5 M), whereas the salt-out strategy relies on the accumulation of compatible solutes to counteract the external osmotic pressure caused by high salinity.^11^ This creates two distinct intracellular conditions to which the proteome must adapt. Therefore, salt-in organisms might show acidic residue enrichment and lower pI compared with salt-out organisms, creating further compositional diversity within halophiles.

Although proteome-wide compositional signatures associated with temperature and salinity have been reported, psychrophilic adaptation has received less systematic attention. Comparisons including thermophiles, psychrophiles, halophiles, and mesophiles within a common framework remain limited.^4,12^ Such compositional signatures could inform comparative proteomics, protein evolution, and environmental microbiology. They can also assist in identifying extremophiles from environmental sequence data.^13^

Here, we compared proteome-wide compositional patterns across thermophiles, psychrophiles, halophiles, and mesophiles. We analyzed 40 complete proteomes to measure the mean protein length, mean % acidic residues, mean % basic residues, acidic-to-basic ratio, mean protein grand average of hydrophobicity, and mean predicted pI. Furthermore, we used principal component analysis to assess whether the four groups occupy distinguishable regions in the proteome compositional space. We tested whether adaptation to high salinity, high temperature, and low temperature is associated with proteome-wide compositional differences relative to mesophiles. Based on previous studies, we expected halophiles, particularly salt-in species, to exhibit a higher percentage of acidic residues and a lower pI, thermophiles to exhibit differences in hydrophobicity and charged residues, and psychrophiles to exhibit characteristics similar to mesophiles at this resolution.^8,9,10^

## METHODS

### Organism Selection

Organisms were assigned to four environmental groups based on their optimal growth temperatures or their salinity requirements. The groups included thermophiles (optimal growth temperatures 50 to 100°C), halophiles (optimal salinity 1 to 30%), psychrophiles (cold-adapted organisms with optimum growth temperatures -5 to 25°C), and mesophiles (optimal growth temperatures 25 to 37°C), with mesophiles serving as the comparison group. A total of 40 organisms were selected, with 10 organisms representing each environmental group. The groups comprised of eight bacterial and two archaeal species for prokaryotic inclusivity. Eukaryotes were excluded. Complete proteomes were downloaded in FASTA format from UniProt release 2026_02 (access date 10/07/2026). The species, strains, environmental groups, and UniProt proteome accession numbers of all selected organisms are given in Table 1.

**Table 1.**
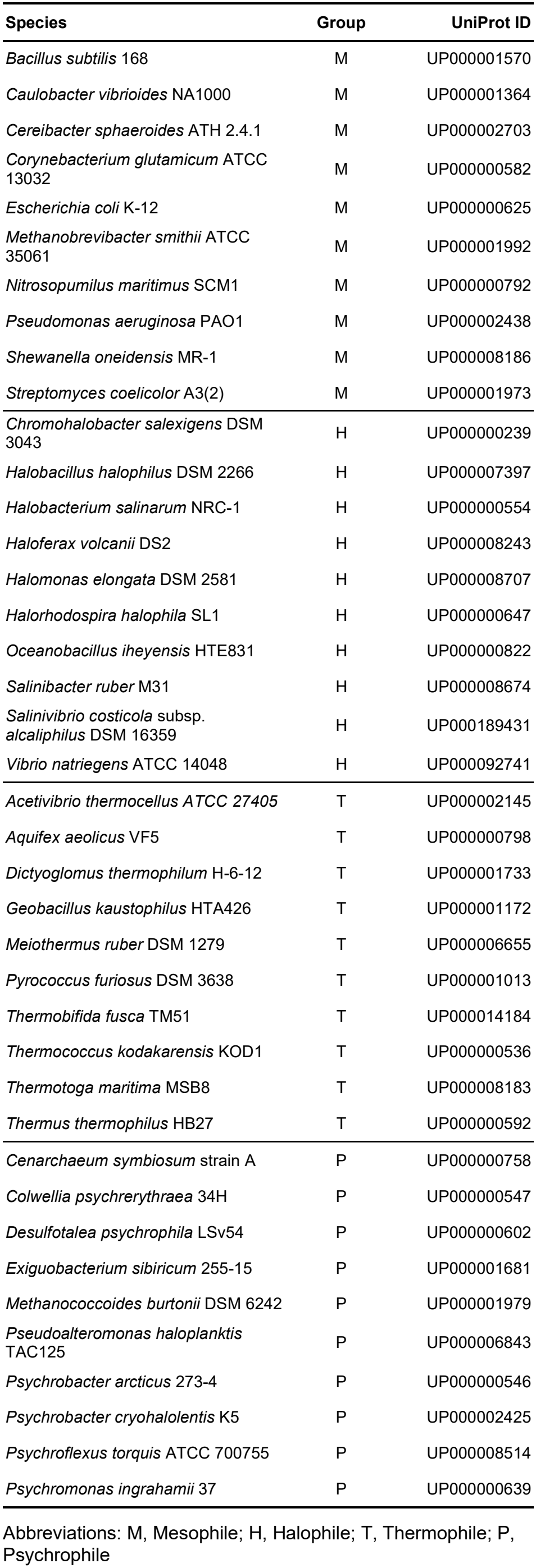
Organisms and UniProt proteome IDs included in the analysis.

### Proteome Processing and Calculation of Proteome-Wide Metrics

Proteome sequences were processed, and the proteomic metrics were calculated in R version 4.5.1. The Biostrings, tidyverse, Peptides, and writexl packages were used to import amino acid sequences, organize and summarize data, calculate predicted isoelectric points, and export the processed datasets, respectively.

Hydrophobicity was calculated using the Kyte-Doolittle scale. The hydropathicity values assigned to the amino acid residues in each protein sequence were then averaged to obtain a single grand average of hydropathicity (GRAVY) value. Predicted isoelectric points were calculated at the protein level by supplying each protein sequence to the Peptides::pI() function.

Next, protein length, percentage of acidic residues, percentage of basic residues, acidic-to-basic ratio, GRAVY, and predicted pI were calculated for each protein and subsequently averaged across all proteins within each organism’s proteome to obtain an organism-level mean for each metric. Protein-level values were aggregated using an unweighted arithmetic mean so that each protein contributed equally to the organism-level mean regardless of sequence length. Only the 20 canonical amino acids were included in the analysis. Acidic residues were defined as aspartate and glutamate, whereas basic residues were defined as arginine, lysine, and histidine. Proteins containing no basic residues were assigned a missing acidic-to-basic ratio.

Finally, the organism-level datasets were exported to an Excel workbook.

### Statistical Analysis and Data Visualization

All statistical analyses were performed using GraphPad Prism (version 9.5.1). For each of the six proteome metrics (mean protein length, percentage of acidic residues, percentage of basic residues, acidic/base ratio, mean hydrophobicity (GRAVY), and mean predicted pI), the four environmental groups were compared using a one-way ANOVA with n=10 organisms per group and mesophiles acting as the control.

Two assumptions were made for each metric. Normal distribution was tested using the Shapiro-Wilk test, and homogeneity of variances was tested using the Brown-Forsythe test. When normality was not the case (Shapiro-Wilk P < 0.05), a non-parametric Kruskal-Wallis test was used. However, in the case where normality was maintained, the parametric ANOVA remained. ANOVA test pairwise differences were examined using Tukey’s post hoc test. Conversely, Kruskal-Wallis tests were examined using Dunn’s post-hoc test. η^2^ (equivalent to R^2^ of ANOVA) was used to quantify the proportion of variance in each metric attributed to the said group (effect size), alongside p-values. The six metrics were tested independently; thus, a Bonferroni significance threshold test was used (0.05 ÷ 6 = 0.0083). This was in addition to the conventional threshold (0.05).

Further analysis of the halophile group was performed. Halophiles were split into salt-in and salt-out according to each organism’s given strategy. Group comparisons were repeated with this split in place to assess whether adaptation accounted for group differences.

All figures were created in R version 4.5.1 using the ggplot2 package, while the ggrepel package was used to minimize overlap between the selected species labels. The organism-level dataset was used to visualize variation in six proteome-wide metrics among the four environmental groups. These metrics comprised mean acidic residue percentage, mean basic residue percentage, mean acidic-to-basic ratio, mean protein GRAVY, mean predicted pI, and mean protein length. Each metric was displayed in a separate boxplot panel with the environmental groups labeled on the x-axis. Boxplots were used to summarize the distribution of organism-level values within each environmental group. The boxplots for the six different metrics were brought together to create one comprehensive figure.

A principal component analysis (PCA) plot was created using the prcomp() function to visualize compositional similarity among 40 organisms. It was created using the organism-level dataset and the mean percentage of acidic residues, mean percentage of basic residues, mean protein GRAVY, mean predicted pI, and mean protein length. The acidic-to-basic ratio was excluded to avoid redundancy, as it was derived from the acidic and basic residue percentages. The variables were scaled and centered before PCA. PC1 represented the linear combination of variables that explained the greatest proportion of variance (42%) among organisms. PC2 explained the next greatest proportion of variance (31%) and was orthogonal to PC1. Each organism was assigned a score for both PC1 and PC2, which determined its position on the x and y axes of the PCA plot, respectively. The percentages on the axes represented the total proportion of variance, ellipses represented the spread of each environmental group in the compositional space, and the arrows represented the PCA loading vectors. The relative arrow length indicated the strength of contribution, whereas the direction indicated the association of each metric with PC1 and PC2.

In all figures, each point represented one organism, and each color represented its respective environmental group. Green represented mesophiles, red represented thermophiles, pink represented halophiles, and blue represented psychrophiles. Pink circles were used only to highlight the three salt-in halophiles (*Haloferax volcanii* DS2, *Halobacterium salinarium*NRC-1, and *Salinibacter ruber* M31). All figures were exported as PNG files.

## Results

### Testing Assumptions

Before groups were compared, two assumptions were carried out. Residuals were normally distributed for four of the six metrics tested (Shapiro-Wilk value P > 0.05). Two out of the six metrics failed: mean predicted pI (P = 0.0028) and acidic-to-basic ratio (P < 0.0001). Therefore, these two metrics used the Kruskal-Wallis test, which subsequently does not assume normality. The remaining four used the one-way ANOVA test. Said groups showed homogeneous spread (Brown-Forsythe P > 0.15), so ordinary ANOVA tests were used.

### Proteome charge differed significantly between extremophile groups

Out of the six metrics, four differed significantly. These groups all passed a stricter Bonferroni threshold test of P < 0.0083 with each metric being a measure of protein charge.

### Predicted pI

The largest difference was observed in mean predicted pI, a measure of the proteome’s acidity/basicity (Kruskal-Wallis P value = 0.0001). Group membership accounted for about half the variation (η^2^ = 0.505). Halophiles were the most acidic (pI 6.09 ± 0.62), differing from thermophiles (7.37 ± 0.39, P < 0.0001), psychrophiles (6.87 ± 0.43, P = 0.0044), and mesophiles (6.87 ± 0.44, P = 0.0042). The other pairs did not differ significantly. This suggests that halophiles may show a distinct separation rather than a gradual transition across the four groups (Figure 1E).

**Figure 1.**
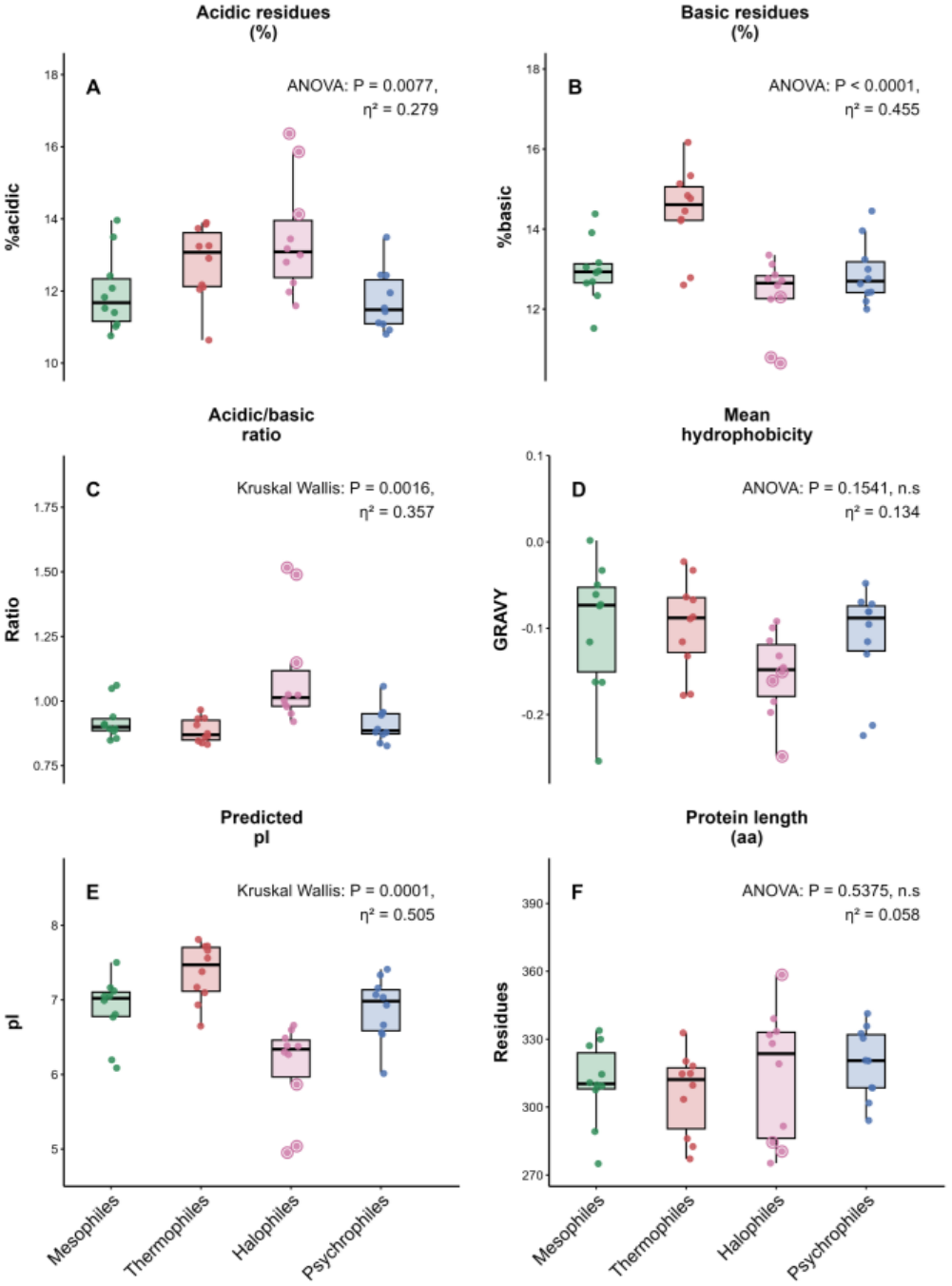
Proteome-wide metrics across environmental groups labeled A-F. Each data point represents one organism (n = 10 per group). Horizontal bars indicate group means and vertical bars show ± 1 standard deviation. Points outlined in pink circles are the three salt in halophiles: Halobacterium salinarum NRC-1, Haloferax volcanii DS2, Salinibacter ruber M31. Test statistics and effect sizes are shown in each panel (n.s = not significant).

### Basic Residues

The percentage of basic residues (K + R + H) showed the second largest separation (F = 10.02, P < 0.0001, η^2^ = 0.455). Thermophiles were the outlier, containing more basic residues (14.45 ± 1.09%) than halophiles (12.34 ± 0.92%, P < 0.0001), psychrophiles (12.91 ± 0.78%, P = 0.0027) and mesophiles (12.96 ± 0.79%, P= 0.0038). All three latter groups did not differ from one another (all P > 0.43) (Figure 1B).

### Acidic Residues

The percentage of acidic residues (D + E) showed the weakest difference of the four-charge metrics (F = 4.638, P = 0.0077, η^2^ = 0.279), narrowly clearing the Bonferroni threshold under ANOVA. Halophiles (13.46 ± 1.58%) exceeded psychrophiles (11.72 ± 0.86%, P = 0.0105) and mesophiles (11.96 ± 1.07%, P = 0.0329) but did not differ from thermophiles (P = 0.5785).

### Acidic-to-Basic Ratio

The acidic-to-basic ratio followed the halophile distinction (Kruskal-Wallis P value = 0.0016, η^2^ = 0.357). Halophiles showed an elevated ratio (1.10 ± 0.22) greater than thermophiles (0.89 ± 0.05, P = 0.0017), psychrophiles (0.91 ± 0.07, P = 0.0059), and mesophiles (0.92 ± 0.07; P = 0.0116).

### Protein length and hydrophobicity showed no group differences

In contrast to charge metrics, neither protein length (F = 0.736, P = 0.5375, η^2^ = 0.058) nor GRAVY (F = 1.859, P = 0.1541, η^2^ = 0.134) differed between groups. Both results were subsequently confirmed by Kruskal Wallis tests (P value = 0.5057 and P value = 0.1073). The average protein length differed by fewer than 14 amino acids across all four groups, and average GRAVY had a range of −0.15 to −0.10. Thus, thermophile proteins were not more hydrophobic than mesophiles, and halophile proteins were not shorter (Figure 1D and 1F).

### Psychrophile proteomes showed no differences from the control

Psychrophiles did not differ significantly from mesophiles on any of the six metrics. These comparisons produced the highest P values in the dataset (pI = 1.0000, basic residues P = 0.9994, acidic/basic ratio P = 0.9939, and acidic residues P = 0.9688). No proteome-wide signature for cold adaptation was therefore detected at this level of resolution.

### Principal Component Analysis (PCA)

PCA reduced five of the six metrics into two axes. These axes accounted for 73.8% of the total variance in the data. PC1 (42.5%) was driven mainly by pI and % acidic residue content, while PC2 (31.3%) was comprised mainly by % of basic residues and protein length. Leaving charge as the main axis of variation in the figure. In the resulting plot (Figure 2), thermophiles separated along the % of basic residue axis. Psychrophiles completely overlapped with mesophiles. Halophiles were the most widely spread, with three halophile organisms sitting well apart from the rest of the dataset.

**Figure 2.**
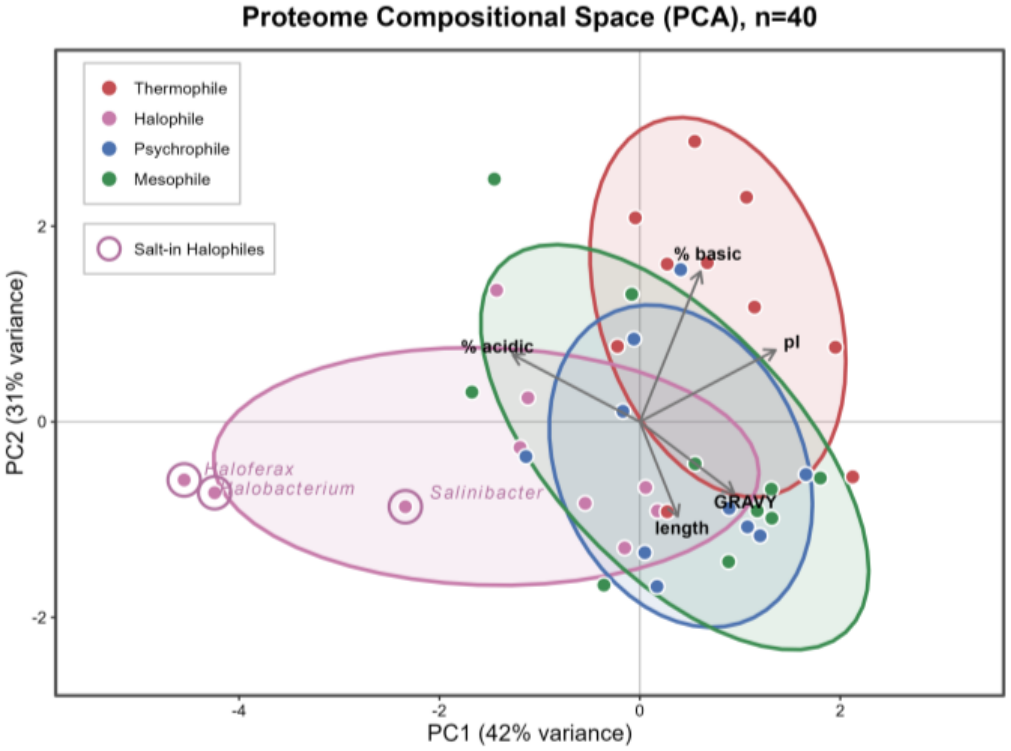
Principal component analysis (PCA) of proteome composition. The variable loadings for PCA are represented as arrows excluding the acidic-to-basic ratio to avoid redundancy. The three salt-in halophiles labelled lie outside the range of all other groups. Psychrophiles and mesophiles overlap almost completely.

### Halophile proteomes are separated by Salt strategy (exploratory analysis)

The three outlying halophiles identified by PCA were Halobacterium salinarum NRC-1, Haloferax volcanii DS2, and Salinibacter ruber M31. All three were salt-in organisms, with the halophile acidic signal being confined to these individuals. Their mean pI values were 4.95, 5.04, and 5.87, respectively. The remaining seven salt-out halophiles, however, lied within a narrower range (6.27-6.66), subsequently overlapping with the controls. Furthermore, the mean acidic-to-basic ratio was higher in salt-in organisms, 1.385, compared to salt-out organisms, 0.984 and 0.924 in mesophile controls.

Repeating the comparison with halophiles divided into these two subgroups based on salt strategy increased the variation for every charge metric. Effect size (η^2^) rose for the acidic-to-basic ratio (0.357 to 0.758), % acidic residues (0.279 to 0.529), pI (0.505 to 0.674), and % basic residues (0.455 to 0.550) (all five-group P ≤ 0.00002). Halophiles still differed slightly from mesophiles in terms of pI (P = 0.0131) and acidic-to-basic ratio (P = 0.0458). Though this effect was not significant in % acidic residue content (P = 0.1493).

This indicates a gradient between the two adaptation strategies rather than a clear separation (Figure 3).

**Figure 3.**
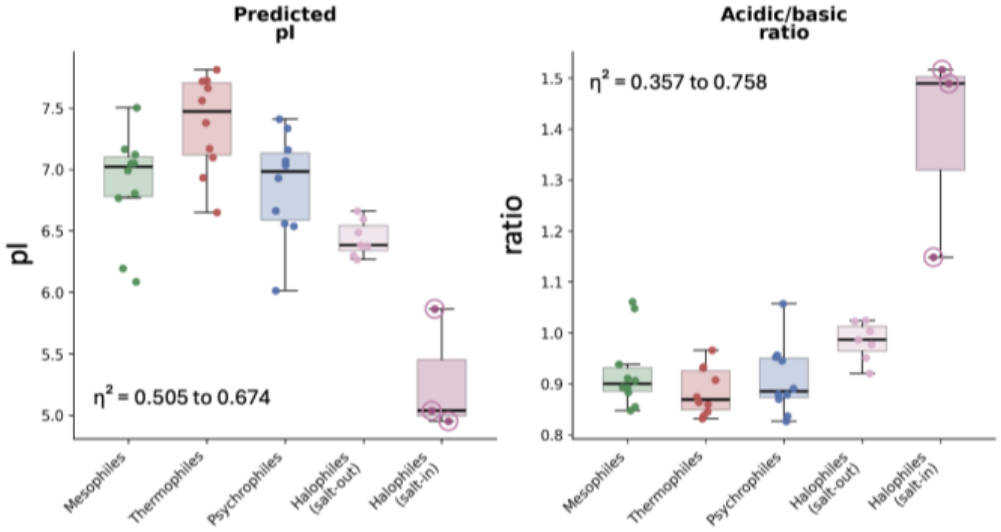
Predicted pI and Acidic/basic ratio graphs with halophiles separated into salt-in (n = 3) and salt-out (n = 7) subgroups. Salt-in organisms are circled in pink. Effect size (η^2^) values shown in the corners of their respective subtypes.

## DISCUSSION

Charge-related properties were the clearest proteome-wide signatures across the four environmental groups. Halophiles exhibited a distinctly low mean predicted pI as well as an elevated acidic-to-basic ratio, whereas thermophiles exhibited an increased mean basic residue percentage. The acidic residue percentage differed across the four groups, with halophiles showing higher values than psychrophiles and mesophiles. However, the pairwise differences did not remain significant under the Bonferroni threshold. In contrast to the charge-related properties, mean protein length and mean GRAVY did not differ significantly. Psychrophiles also did not differ significantly from mesophiles and showed extensive overlap in the PCA. These results broadly supported our hypotheses on the acidic characteristics of halophiles, basic residue enrichment of thermophiles, and the compositional similarities between psychrophiles and mesophiles. An exception to our hypotheses was thermophiles not exhibiting differences in mean hydrophobicity.

On average, halophiles exhibited greater acidic characteristics, including a lower mean predicted pI value and an elevated mean acidic-to-basic ratio, supporting previous findings for this group.^4,8,9^ However, the three PCA outliers were all salt-in halophiles, and the effect size was stronger when salt-in and salt-out organisms were analyzed separately. Salt-out organisms had pIs that overlapped with mesophiles and were closer to mesophiles in their acidic-to-basic ratios, whereas salt-in organisms had distinctly lower pIs and a higher acidic-to-basic ratio. This suggests that the acidic signature mainly belongs to the salt-in subgroup. Furthermore, it is argued that the high negative surface charge of halophilic proteins prevents aggregation and renders them more soluble and flexible at high salt concentrations, explaining the characteristics observed in the salt-in organisms.^14^ The overlap between salt-out organisms and mesophiles may indicate that compatible solute-based osmoregulation imposes weaker proteome-wide compositional constraints than adapting to high intracellular salt concentrations.^10^ However, this explanation remains speculative.

Thermophiles exhibited comparatively high charged residue percentages of both signs, which supported the initial hypothesis. This is associated with the need for increased protein stability under extreme heat via the formation of salt bridges.^8^ Conversely, the mean GRAVY value did not demonstrate any significant difference at this resolution.

These findings do not necessarily rule out hydrophobic interactions for thermophiles since a proteome-wide mean cannot account for local hydrophobic core packing.Furthermore, organism-level GRAVY was calculated as an unweighted mean of protein-level GRAVY scores. The result therefore reflected mean protein hydropathicity rather than the residue-weighted hydropathicity of the proteome. Overall, the detected thermophilic signature was associated with the charged residue percentages rather than the GRAVY scores.

Psychrophiles did not differ from mesophiles on any of the six metrics and broadly overlapped in the PCA. The five variables included in the PCA did not distinguish between the two groups. This finding was consistent with our hypothesis across the metrics examined. However, the metrical similarities should not be taken as evidence for a lack of proteomic adaptation in psychrophiles. Cold adaptation includes shifts in individual amino acids both across the proteome and in secondary structural elements.^10^ Therefore, it is possible that these signatures were obscured when residues were grouped into broader metrics. It is also possible that the PCA missed the signals as the principal components were driven mostly by charge-related variables.

Additionally, the lack of any psychrophilic signatures may also be explained by the relatively small sample size. An increased sample size could improve precision, as only 10 organisms represented each group. The conclusions about salt-in organisms were particularly uncertain because the group contained only three organisms. Furthermore, organisms were selected using predefined criteria rather than at random, which might have introduced a sampling bias. The analyses did not account for phylogenetic non-independence, meaning that similarities among closely related organisms may have reflected shared ancestry rather than environmental adaptation. Moreover, eukaryotes were excluded, which may limit the generalization of the findings to eukaryotic extremophiles.

The analysis was done using complete UniProt proteomes, which represent the encoded protein repertoire rather than condition-specific protein abundance or expression. Consequently, the calculated proteome-wide composition may not reflect the subset of proteins expressed under the environmental conditions examined. Additionally, an unweighted averaging approach was used for all the metrics, meaning that long proteins did not contribute more than short proteins. Only six metrics were analyzed, which might have obscured variation in individual amino acid frequencies, protein families, or local structural regions. Among these, GRAVY and predicted pI values were computationally calculated rather than experimentally validated.

Future work could increase the sample size, especially for independently selected salt-in and salt-out organisms. Subsequent studies could also use phylogenetically balanced samples or phylogenetic comparative models. Furthermore, the frequencies of all 20 amino acids could be analyzed to determine any signature at that level. Further research may also include the aliphatic index, flexibility-related metrics, and structural features in addition to the six metrics that were analyzed. Additionally, comparing the unweighted protein means with residue-weighted whole proteome calculations could determine whether the group differences were robust to the aggregation method or influenced by variation in protein length. Finally, the identified compositional signatures could be tested on an independent set of proteomes to evaluate their utility for identifying extremophiles. Only after independent validation should the metrics be developed into a classification method to identify extremophile candidates from environmental sequence data.

## CONCLUSION

This study asked whether extremophile adaptations leave a measurable compositional signature in an organism’s proteome. Our findings supported the presence of such signatures along the charge axes. Thermophiles were enriched in basic residues, while salt-in halophiles were strongly acidic. Protein length and hydrophobicity carried no detectable signal across the groups. Subsequently, the hypothesis of thermophilic proteins being more hydrophobic was not supported in our findings.

A possible finding was that halophiles are not a coherent proteomic category. Subdividing halophiles by salt strategy revealed that the acidic signal was confined to salt-in organisms while salt-out organisms resembled mesophiles more closely. These results indicated that survival strategy is a more important predictor of proteome composition than the environmental group.

No psychrophilic signature was detected. This is consistent with the idea that cold-adapted organisms operate through localized structural flexibility. However, it may equally reflect the limitations of the metrics used. Such signatures might potentially arise if other metrics tested for structure rather than composition alone.

Compositional fingerprinting could potentially identify extremophiles from sequence alone, only if the adaptations are charge-based. This same logic may apply to tumours that adapt to acidic and hypoxic microenvironments. However, it is hard to speculate without similar compositional studies for eukaryotic proteomes. Future work should sample within strategy-defined subgroups and extend analysis to structural features to capture cold adaptations.

## Supporting information

Proteome Metrics for All Organisms

## AUTHOR INFORMATION

Corresponding Authors

## Author Contributions

Kayra Güvenç Duran conceived and designed the study, developed the R analysis pipeline, performed the proteome-wide computational analysis, generated the figures and led manuscript writing.

Mahamed Salah curated and verified the halophile and psychrophile proteome datasets. Performed all statistical analysis and data visualisation, and drafted the Results, Abstract and Conclusion.

## Abbreviations

GRAVY: Grand Average of Hydrophaticity
PCA: Principal Component Analysis

